# A simple network of proprioceptive reflexes can produce a variety of bipedal gaits

**DOI:** 10.1101/2025.02.05.636668

**Authors:** Elsa K. Bunz, Daniel F. B. Haeufle, Syn Schmitt, Thomas Geijtenbeek

**Affiliations:** Institute for Modelling and Simulation of Biomechanical Systems, University of Stuttgart, Stuttgart, Germany; Stuttgart Center for Simulation Science, University of Stuttgart, Stuttgart, Germany; Center for Bionic Intelligence Tuebingen Stuttgart, Tuebingen Stuttgart, Germany; Institute of Computer Engineering, Heidelberg University, Heidelberg, Germany; Hertie Institute for Clinical Brain Research and Center for Integrative Neuroscience, Tuebingen, Germany; Goatstream, Utrecht, the Netherlands

## Abstract

Proprioception is crucial for movement, yet the role of proprioceptive reflexes in legged locomotion is still poorly understood. While previous simulation studies have shown great potential for reflex-based control strategies, these controllers are typically catered to specific gaits, using hand-crafted feedback pathways that are linked to specific gait phases. In this work, we explore the control capabilities of a simple reflex controller that consists of only monosynaptic and antagonistic length and force feedback pathways with constant gains. Despite its simplicity, we found our control framework capable of producing a wide variety of natural gaits, including walking and hopping, forwards and backwards, and running in different variations and at different velocities – without requiring any rhythmic inputs or high-level state machines modulating the feedback gains. Our work highlights the important role and flexibility of proprioceptive reflexes and suggests a necessary re-evaluation of their role in locomotion. Due to its simplicity and flexibility, our control framework provides a solid basis for the development of higher level neuromuscular control systems.

**Author summary:** Human locomotion is characterized by an impressive versatility and humans transition naturally between different gaits. However, the neural mechanisms underlying human motor control are not fully understood and remain controversial. The two widely accepted basic components of the neural circuitry involved in locomotion are central pattern generators (CPG) and spinal reflexes, leading to a long-standing debate on their respective roles. Neuromusculoskeletal models can be employed to study the role of these neural control structures in simulation. Until now, these models mostly focus on walking and require an additional component like a state-machine or CPG besides reflexes to generate rhythmic activation of muscles. In this work, we propose a controller that can generate different rhythmic gaits like walking and hopping, forwards and backwards, as well as running, based solely on simple reflexes known to exist in the mammalian spinal cord. We challenge the current view of the role of rhythm-generating components and show the remarkable potential of reflexes for locomotion control. To our knowledge, this is the first work showing that versatile locomotion can be achieved solely based on reflexes without the need of a CPG. Our results contribute to the understanding of human motor control and are relevant for the control of robots, exoskeletons, prostheses and ortheses.

## Introduction

The role of proprioceptive reflexes in human locomotion has been subject to longstanding research and debate [1, 2]. Neuroscientific research often emphasizes the importance of central pattern generators (CPGs), i.e. spinal circuits that can produce rhythmic activity in the absence of afferent or supraspinal inputs [3, 4]. Yet, information from proprioceptive sensors is indispensable for movement generation [5] and has been shown to be essential for the regulation of stepping [6] and the modulation of CPG output [7, 8]. However, the way in which CPGs and reflexes functionally interact is poorly understood, and in vivo preparations have not yet reached the level of detail to decipher parameters and structures related to CPGs and reflexes, especially in dynamic locomotion [3].

Predictive neuromuscular simulations are a useful alternative to in vivo preparations, since they allow complete and fine-grained control over the experimental setup, and help identifying the potential role of individual control primitives [1, 9]. In addition, predictive neuromuscular simulations have important potential applications, such as aiding in the design of assistive devices through human-in-the-loop simulation [10–14], predicting the outcome of medical procedures [15–17], and predicting gait stability and perturbation responses [18–20]. However, to fulfill these promises, a control framework is needed that is both neurologically grounded and capable to replicate the versatility of human movement.

An important simulation study that highlights the potential of reflexes in locomotion is the work of Geyer & Herr [21], who developed a locomotion control strategy that is driven mostly by proprioceptive reflexes. Their work inspired a vast number of derivative controllers including extensions to 3D and the addition of higher-level control components [22–26] and was the basis for several subsequent predictive simulation studies on human gait [27–29]. All these controllers produce rhythmic muscle activation by switching between task specific states (e.g. swing and stance). This hand-crafted state selection logic activates and deactivates specific reflex pathways based on the current state of each leg (i.e. based on ground contact). Furthermore, the reflex pathways are domain-specific, as they are manually picked to mimic human level-ground walking kinematics and muscular activations. This tailoring to level-ground walking and dependence on state-switching limits their ability to generalize to other gaits. Works extending the controller of [21] to other environments or movements mostly focus on adding a higher-level component [18, 23] ([22] also adds new reflex pathways in addition to a supraspinal layer) or on further refining and tailoring the used states [25, 26]. While all of these works are valuable and study different parts of the human motor system this approach likely obscures the potential within reflexes themselves. The next step in motor control research extending from these works is to look for complex tasks [30] and a solid spinal basis is necessary to approach this.

The purpose of this study is therefore to explore the locomotor capabilities of proprioceptive reflexes in the absence of a high-level state selection or modulation. Instead, we use delayed length and force reflex pathways with constant feedback gains which are active throughout the simulation, relying solely on reciprocal innervation to produce alternating rhythmic activity. Thereby, we devise a more general network of reflexes to allow generalization to other gaits besides walking. For the reflex connectivity, we mimic the basic organization found in the mammalian spinal cord [7] by allowing reflex pathways to homonymous and antagonistic muscles.

In spite of its simplicity, we found that these basic reflexes were sufficient to produce a variety of natural bipedal gaits like walking, hopping and running in different variations and at different velocities – only by selecting different sets of constant feedback parameters. Even though our controller still misses many components present in human motor control, its simplicity allows to study the remarkable potential of reflexes. In contrast to current beliefs [31], our results suggest that the simple proprioceptive reflexes found in the spinal cord are alone capable of producing the basic rhythmic activity required for locomotion, independent from CPGs.

## Materials and methods

### Model

Simulations were performed using a planar nine degree-of-freedom model representing an adult human male of 74.5 kg and 1.8 m height. The model consists of seven segments (torso and two legs with femur, tibia, foot). Hip, knee and ankle joint of each leg are actuated by 9 Hill-type musculotendon units with elastic tendons and muscle fiber damping [32]: gluteus (GLU), iliopsoas (ILI), rectus femoris (RF), hamstrings (HAM), biceps femoris short head (BF), vastus (VAS), gastrocnemius (GAS), tibialis anterior (TA) and soleus (SOL) (see also Fig. 1). Both the mass properties and muscle parameters in our model (optimal fiber length, tendon slack length, pennation at optimal fiber length, maximum isometric force) were taken from [33], with the exception of the hamstring parameters, for which we used updated data from [34]. We further adapted the via points of the muscle pathways to match moment arm data from [35], which uses wrapping geometry to match experimentally measured moment arms. We avoided the use of wrapping geometry in our model for performance reasons, but found that through via points we could match the moment arms within the margin of error of the source material. In cases where we changed the muscle geometry, we adjusted the tendon slack lengths in such a way that the muscle fiber lengths remained optimal at the same joint angle (or joint angles for biarticular muscles), thereby minimizing changes in the angle at which passive muscle force commences. The latter is important because our control strategy relies on force feedback, which is sensitive to the passive muscle force that is triggered after a muscle stretches past its optimal fiber length. Following that same rationale, we adjusted the tendon slack length of the ankle plantarflexors in such a way that these muscles better match passive dorsiflexion muscle force measurements found in [36]. The curves representing tendon stiffness, as well as the force-length and force-velocity relations are taken from [32], including the passive damping (*β* = 0.1), allowing muscle activation levels to be zero. We use a v_max_ of 10 m/s, similar to [33] and [35]. Muscle activation is computed using a first-order dynamics model, with an activation time constant of 0.01 s and a deactivation time constant of 0.04 s [37].

**Fig 1.**
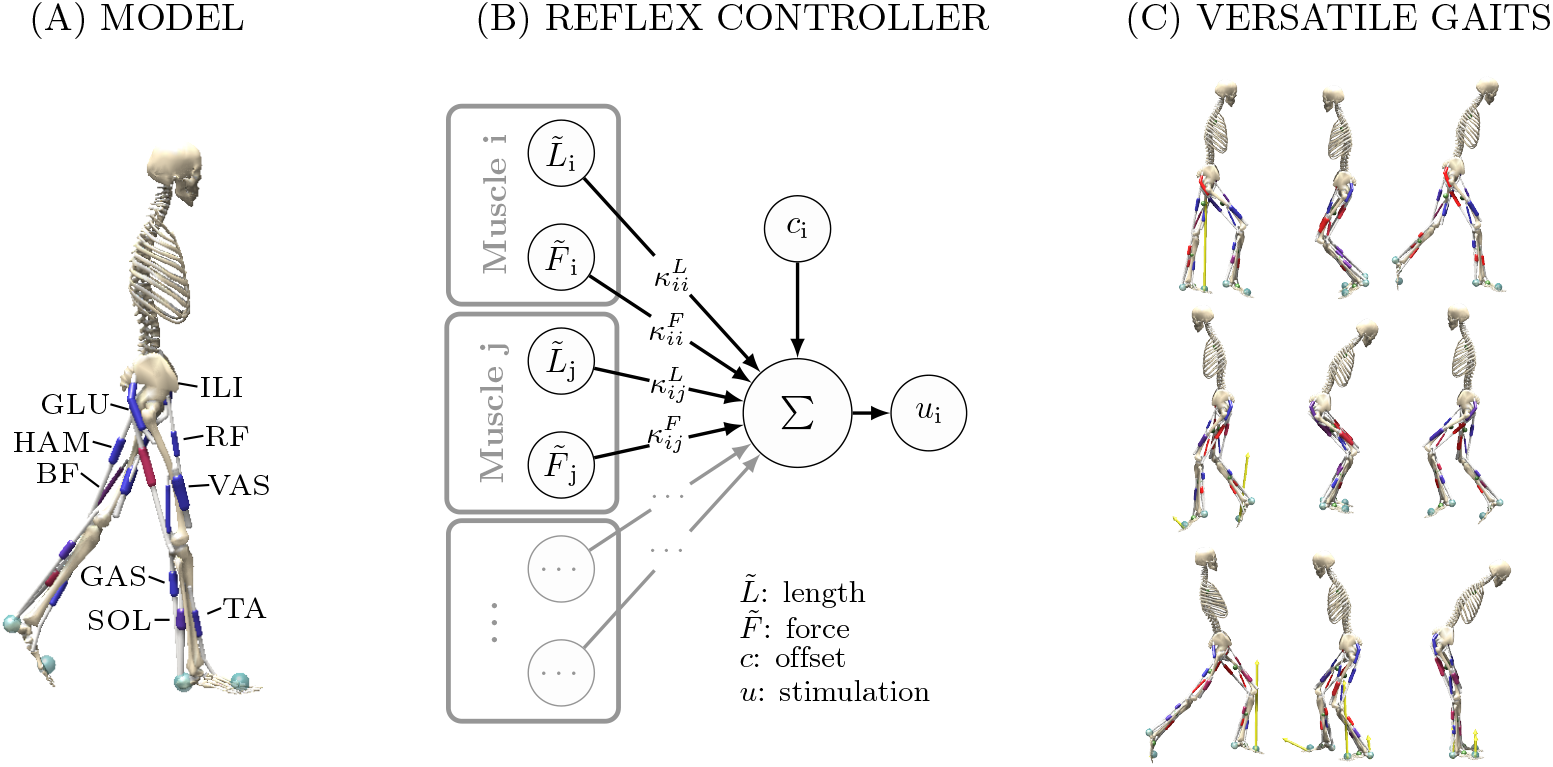
Overview. Our reflex controller can produce a variety of gaits in a human musculoskeletal model. (A) The planar musculoskeletal model consists of seven segments connected by 6 joints which are actuated by 18 Hill-type muscles. (B) Our reflex controller calculates stimulation for each muscle *i* from delayed force 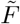 and length 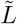 feedback with gains *κ*_*ij*_ and a constant offset *c*_*i*_. Implemented pathways include homonymous and antagonistic pathways for length and force. Feedback gains and offsets are optimized based on high-level objectives. (C) The controller can produce a variety of bipedal gaits like forwards and backwards walking and hopping as well as running.

Ground contacts are modeled using the Hunt-Crossley contact model [38] with two contact spheres (*r* = 0.03, stiffness = 5 10^6^, dissipation = 1) per foot segment, using the friction model implemented in OpenSim [39] (static friction = 0.9, dynamic and viscous friction = 0.6). The model is implemented in SCONE [40] and Hyfydy [41] and is provided as supplementary material. An overview of the model parameters is given in Table S1.

### Controller

Our neural control model is based only on scalar muscle fibre length and muscle tension signals. They represent a common simplification of proprioceptive feedback from muscle spindles (type Ia sensory fibers), and Golgi tendon organs (type Ib sensory fibers), respectively [7, 21]. We acknowledge that muscle spindle output can not be predicted only based in muscle fibre length, since also velocity, force and yank [42] play a role, even though experimental results have not yet been fully explained by any of these variables [43]. For simplicity we therefore focus on the commonly adopted modeling of proprioceptive feedback as muscle fiber length and tension and leave a detailed study and refinement of potential sensor signals for future work.

Following the basic organization found in the mammalian spinal chord [5, 7, 44], our model uses two types of reflex connections: *homonymous* connections to the same muscle and *antagonistic* connections to all antagonist muscles. For Ia length reflexes, homonymous connections are considered monosynaptic and constrained to be excitatory, while antagonistic connections are constrained to be inhibitory (via the Ia inhibitory interneuron). Ib force reflexes are allowed to be either excitatory or inhibitory, since they connect via both inhibitory and excitatory interneurons [7].

For our model with nine muscles per leg this leads to connection matrix containing 9 monosynaptic and 22 antagonistic connections (Fig. 2). All pathways are modeled with neural delays Δ*t* [10 ms, 35 ms] based on experimental data of H-reflex latency as well as the length of the pathway (i.e. the more distal the more delay), following [45] (for the delay values see Table S1).

**Fig 2.**
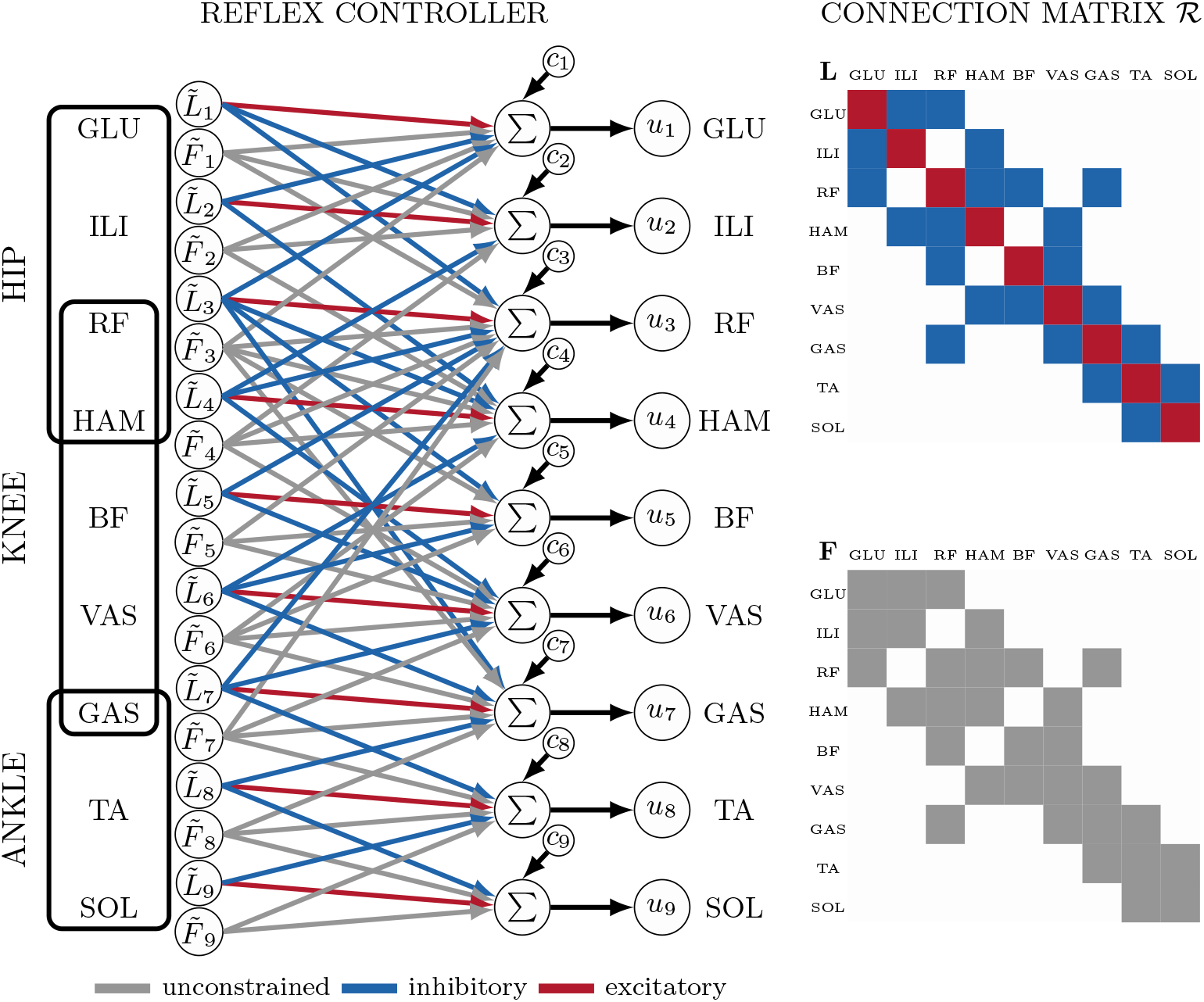
Control architecture. Muscle stimulations *u*_*i*_(*t*) for the nine muscles per leg are computed through delayed length 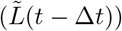 and force 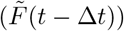 feedback. The connection matrix ℛ (right) contains both, homonymous connections innervating the muscle based on its own sensor signal (red in *L* matrix) and connections between antagonistic muscles (blue in *L* matrix). Gains are either unconstrained (grey, 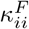 and 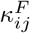), constrained to be excitatory (i.e. ≥ 0, red, 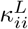) or inhibitory (i.e. ≤ 0, blue, 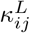). Including the constant thresholds *c*_i_, the reflex controller contains 71 free parameters.

Overall, the controller thus produces a stimulation *u*(*t*) of each muscle *i* based on constant-gain delayed length 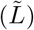 and force 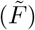 feedback, and a constant offset *c*_*i*_:

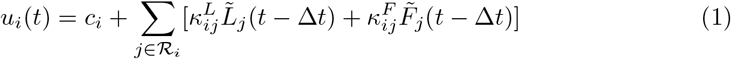

Here, *u*_*i*_ is saturated to [0,1], |*κ*|≤2, |*c*|≤1. We use the same pathways and control parameters (*κ* and *c*) for both legs. Our controller does not include cross-connections between the left and right leg. The normalized muscle fiber length 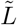 and force 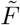 are defined as:

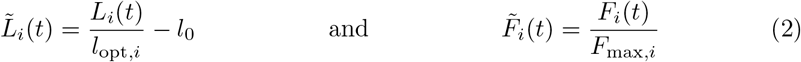

with *l*_opt,*i*_ the optimal fiber length and *F*_max,*i*_ the maximal isometric force of muscle *i* (see Table S1). We set *l*_0_ = 0.5 to ensure 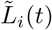 within a good range while avoiding a possible change of sign if *l*_0_ is set too high. All reflex gains and offsets are constants and there is no modulation or switching based on different states.

### Optimization

Our reflex controller contains 71 free parameters (9 *c*_*i*_, 31 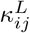 and 31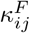), which dictate the behavior produced by the controller. We explore the capabilities of the reflex controller by defining five target gaits: forward walking (Walk Fwd) and hopping (Hop Fwd), backwards walking (Walk Bkw) and hopping (Hop Bkw), and running (Run). For the five target gaits, based on initial gaits we found, we extract and manually tune the initial state of the model (torso and joint angles and velocities) to roughly reflect the target gait (see Fig. 3 and Table S2). To produce a periodic motion the reflex controller generally needs to find an equilibrium where the interplay of reflexes produces cyclically reoccurring states and sensor signals. An initial state close to this motion minimizes the initial correction the controller will have to perform. For this reason, the initial state strongly influences the resulting gait and can be used to guide the optimization towards the intended target gait. We do not optimize for specific motions through tracking objectives, but instead minimize a high-level cost function *J*, which consists of a velocity target and a term for minimizing muscle activation as a measure of muscle fatigue [46]:

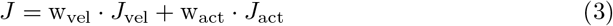

and mainly dictates the direction and speed of the gait. Another included optimization criterion is gait stability for the duration of the simulation *t*_sim_. Simulations are terminated after *t*_max_ = 30 s or if the center of mass (CoM) of the model goes below 0.5 m. We define a gait as stable if it does not fall within the maximum simulation time, i.e. if *t*_sim_ = *t*_max_ and integrate the stability term into *J*_vel_.

**Fig 3.**
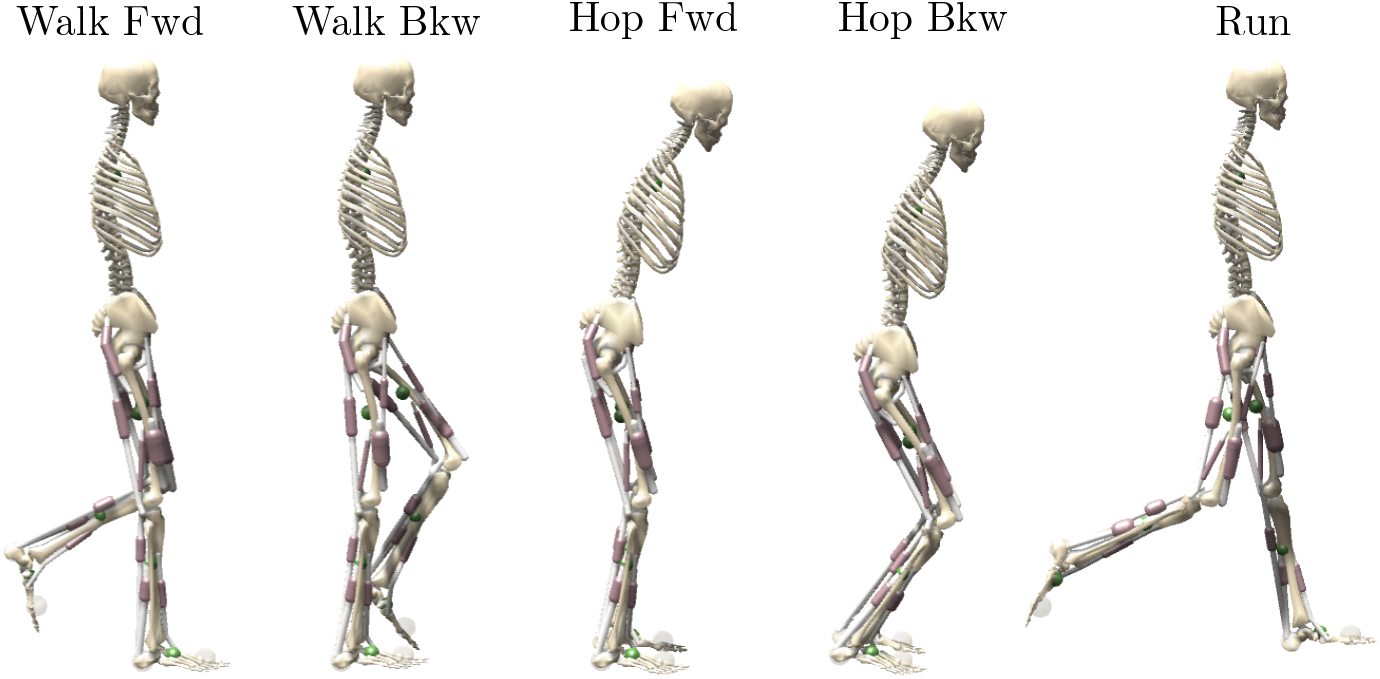
Initial states. For each target gait the simulation starts in a different initial pose (joint angles and velocities) manually tuned to minimize the initial correction the controller has to make and direct the optimization towards the target gait. The numeric values of all initial sets can be found in Table S1.

To optimize for the five different gaits with the same workflow, we furthermore introduce a flag s_v_ into *J*_vel_, which allows to select between the different optimization conditions of reaching a target velocity *v*_tgt_ (flag s_v_ = 0) or maximizing the absolute velocity of the model (flag s_v_ = 1 or 1 for forwards and backwards movements respectively):

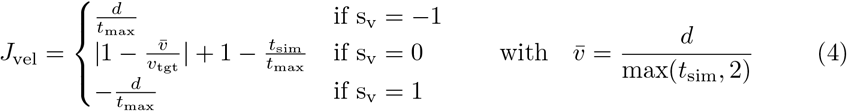

Here, *d* is the completed distance, which corresponds to the horizontal displacement of the CoM of the segment closest to the origin at the end of the simulation (*t*_sim_). *J*_vel_ also encourages stability: if s_v_ ≠ 0, *J*_vel_ is calculated based on the maximum simulation time *t*_max_ (set to 30 s), penalizing early falls where with the same velocity the reached *d* is smaller (and thus *J*_vel_ is smaller) the earlier the model falls. If the optimization target is to reach a target velocity (s_v_ = 0) the average velocity of the model 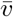 has to be calculated based on the actual simulation time *t*_sim_ to be compared to the target velocity. To foster stable solutions in this case a penalty for early terminations is added 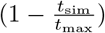. Furthermore, we only calculate the velocity based on *t*_sim_ if *t*_sim_ *>* 2 s as otherwise the optimization gets caught in local minima where the model falls within the first 2 s at a velocity close to the target velocity.

The muscle activation cost term *J*_act_ consists of integrated cubed activation *a* during the time interval 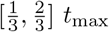s:

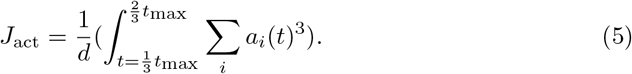

Cubed muscle activation has been suggested as a measure of muscle fatigue [46, 47] based on experimental data indicating a cubic relationship between muscle force and endurance [48, 49]. The restriction to the time interval 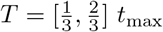helps to initially encourage finding stable gaits before minimizing fatigue 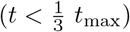 and prevents falling towards the end of the simulation due to micro-optimization of activations 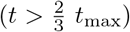. As gains are constant throughout the simulation and the gaits emerge from the interplay of the reflexes an adaptation to minimize the activation during 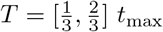 automatically influences the rest of the simulation as well. We do find that the gaits where activation minimization is active show some more inter-stride variability than the ones without, however the strides are not different depending on which time interval they are in (compare Fig. S1a).

With the two cost function terms *J*_vel_ and *J*_act_ we then define the optimization parameters for the five target gaits. Humans typically walk at a preferred walking velocity possibly minimizing energy expenditure [50] and muscle fatigue [46]. The energetically optimal walking speed found by [50] is at 1.23 m/s and the subjects of the experimental dataset we use for comparison show a mean preferred speed of 1.17 m/s [51]. We therefore set a target velocity of 1.2 m/s for forwards as well as backwards walking. Furthermore, we include activation minimization normalized to distance as a cost function term to take muscle fatigue into account [46]. For hopping and running, we optimize for maximum velocity and therefore deactivate the activation cost term. An overview of our optimization parameters can be found in Table 1. As discussed before, with this generic approach the initial state has a large influence on the resulting gait as e.g. hopping forward and running share the same optimization parameters and only differ in their initial state.

**Table 1.**
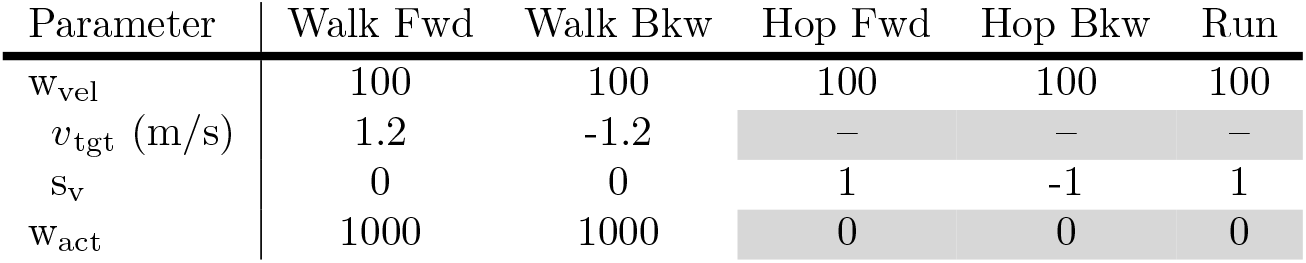
Optimization parameters. Chosen values of the optimization parameters for each of the five target gaits. Cells are greyed out if the cost function term is inactive for the respective gait.

| Parameter | Walk Fwd | Walk Bkw | Hop Fwd | Hop Bkw | Run |
| --- | --- | --- | --- | --- | --- |
| $w_{\text{vel}}$ | 100 | 100 | 100 | 100 | 100 |
| $v_{\text{tgt}}$ (m/s) | 1.2 | -1.2 | — | — | — |
| $s_v$ | 0 | 0 | 1 | -1 | 1 |
| $w_{\text{act}}$ | 1000 | 1000 | 0 | 0 | 0 |

We optimize the parameters using the Covariance Matrix Adaptation Evolution Strategy (CMA-ES) [52]. The initial parameter mean values are rough estimates obtained through experimentation and are provided as part of the supplementary material. To allow sufficient exploration to cover all gaits and avoid local minima, we set the standard deviation to 0.2. We use the same mean values and standard deviations for all optimizations. Optimizations are terminated when the average relative improvement over the last 500 iterations is less than 10^−5^ per iteration.

Due to the complexity and discontinuity of our objective function (which involves a full musculoskeletal simulation in which the model can fall), our optimization problem is subject to many local minima. As a result, the optimization result highly depends on the stochastic sequence drawn by the CMA-ES optimizer. Due to these limitations, we run 20 optimizations for each target gait, each with different randoms seeds (which determine the stochastic sequence used by the optimizer), and pick best result(s). While these results are unlikely to represent the absolute global optimum, we expect them to be close candidates.

### Experimental data

We compare the kinematics and muscular activations of our model to experimental data for forwards walking and running. While humans are capable of performing backwards walking and hopping forward and backwards, unfortunately, there is a lack of experimental data for comparison.

The experimental data of human walking is mostly extracted from [51]. We use the treadmill trials of all participants and extract ten strides at the velocity matching the walking velocity of 1.2 m/s of our model. For each participant, we then average over these strides. Next, we compute the mean and the mean absolute deviation from the mean and display this interval. We also compare to data of one participant, who is closest to our model in terms of weight and height (AB06, 1.8 m, 74.8 kg). As [51] does not contain electromyography (EMG) data for ILI and BF, we furthermore extract data for these two muscles from other sources. ILI is compared to data taken from [53] as used in [21] and BF data comes from [54]. As [53] and [54] do not provide data of multiple subjects, we directly use the extracted data and do not display an interval for ILI and BF. Note, that the EMG data from [51] was normalized for each subject with respect to the average amplitude at a speed of 1.35 m/s, while data for BF from [54] is given in µV and ILI data from [53] was normalized to the maximum manual muscle test value. Therefore, a comparison of magnitude to our simulation data, normalized to maximum activation is difficult.

For running we digitized data from [55] of experienced long distance runners running at 3 m/s. EMG normalization in [55] was performed for each muscle using the maximum voltage across all trials for each subject.

To allow for a quantitative comparison of simulation data with the experimental data, following [21] we compute the maximum cross-correlation *R* of the mean signal and the corresponding time shifts Δ in percent of stride (maximum shift Δ_max_ = 20 %). Again, interpretation of the fit to EMG should be done carefully, as it has been argued that cross-correlation might not be suitable for comparing EMG values (between different subjects) [56].

## Results

For all five target gaits we found stable solutions, i.e. solutions that did not fall within the maximum simulation time (see Video S1). The model reaches the target velocity of 1.2 m/s for forwards and backwards walking and a maximum running velocity of 3.4 m/s. The hopping solutions generate hopping forwards and backwards at a velocity of 1.2 m/s and 1.4 m/s respectively (compare also Table 2).

**Table 2.** Gait velocity. Obtained velocities, stride length and duration for the five target gaits.

|  | Walk Fwd | Walk Bkw | Hop Fwd | Hop Bkw | Run |
| --- | --- | --- | --- | --- | --- |
| Velocity (m/s) | 1.20 | -1.21 | 1.22 | -1.39 | 3.42 |
| Stride length (m) | 1.11 | -0.83 | 0.54 | -0.54 | 1.94 |
| Stride duration (s) | 0.92 | 0.69 | 0.44 | 0.38 | 0.57 |

Our forwards walking gait matches human kinematics well (see Fig. 4, *ϕ*_h,our_: *R* = 0.90, *ϕ*_k,our_: *R* = 0.89, *ϕ*_a,our_: *R* = 0.79). Even though the knee kinematics are not matched as well as by the reflex controller of [21](*ϕ*_k,GH_: *R* = 0.98), the ankle kinematics show a greater maximumn cross-correlation than the model of [21](*ϕ*_a,GH_: *R* = 0.69). The mismatch in knee kinematics is mostly coming from the leg being straight during stance. The vertical ground reaction force shows the characteristic two peaks found in experimental data (albeit with shifted timings, *R* = 0.93). Generally, the computed time shifts Δ of the maximum cross-correlation for knee, ankle and GRF are higher for our model than for the model of [21], but within maximally 3.6 % (Δ_GRF_). All maximum cross-correlation values and times shifts are given in Table 3.

**Table 3.** Maximum cross-correlation. *R* **and time shift** Δ **for joint angles and GRF** Similarity metrics computed between mean of experimental data (exp) or mean data of subject AB06 (for Walk Fwd only) and simulation data from our model (our) or the model of Geyer & Herr [21] (GH, Walk Fwd only).

| | $\phi_h$ | | $\phi_k$ | | $\phi_a$ | | GRF | |
| --- | --- | --- | --- | --- | --- | --- | --- | --- |
| | $R$ | $\Delta$ (%) | $R$ | $\Delta$ (%) | $R$ | $\Delta$ (%) | $R$ | $\Delta$ (%) |
| Walk Fwd |  |  |  |  |  |  |  |  |
| exp-our | 0.90 | 0.0 | 0.89 | 2.1 | 0.79 | 3.4 | 0.93 | 3.6 |
| exp-GH | 0.93 | 0.0 | 0.98 | 0.6 | 0.69 | 1.0 | 0.99 | 0.7 |
| AB06-our | 0.91 | 0.0 | 0.95 | 2.8 | 0.72 | 4.1 | 0.94 | 2.7 |
| AB06-GH | 0.81 | 0.0 | 0.98 | 0.3 | 0.75 | 1.7 | 0.98 | 0.7 |
| Run |  |  |  |  |  |  |  |  |
| exp-our | 0.91 | 0.0 | 0.93 | 2.0 | 0.25 | 20 | 0.78 | 8.5 |

**Fig 4.**
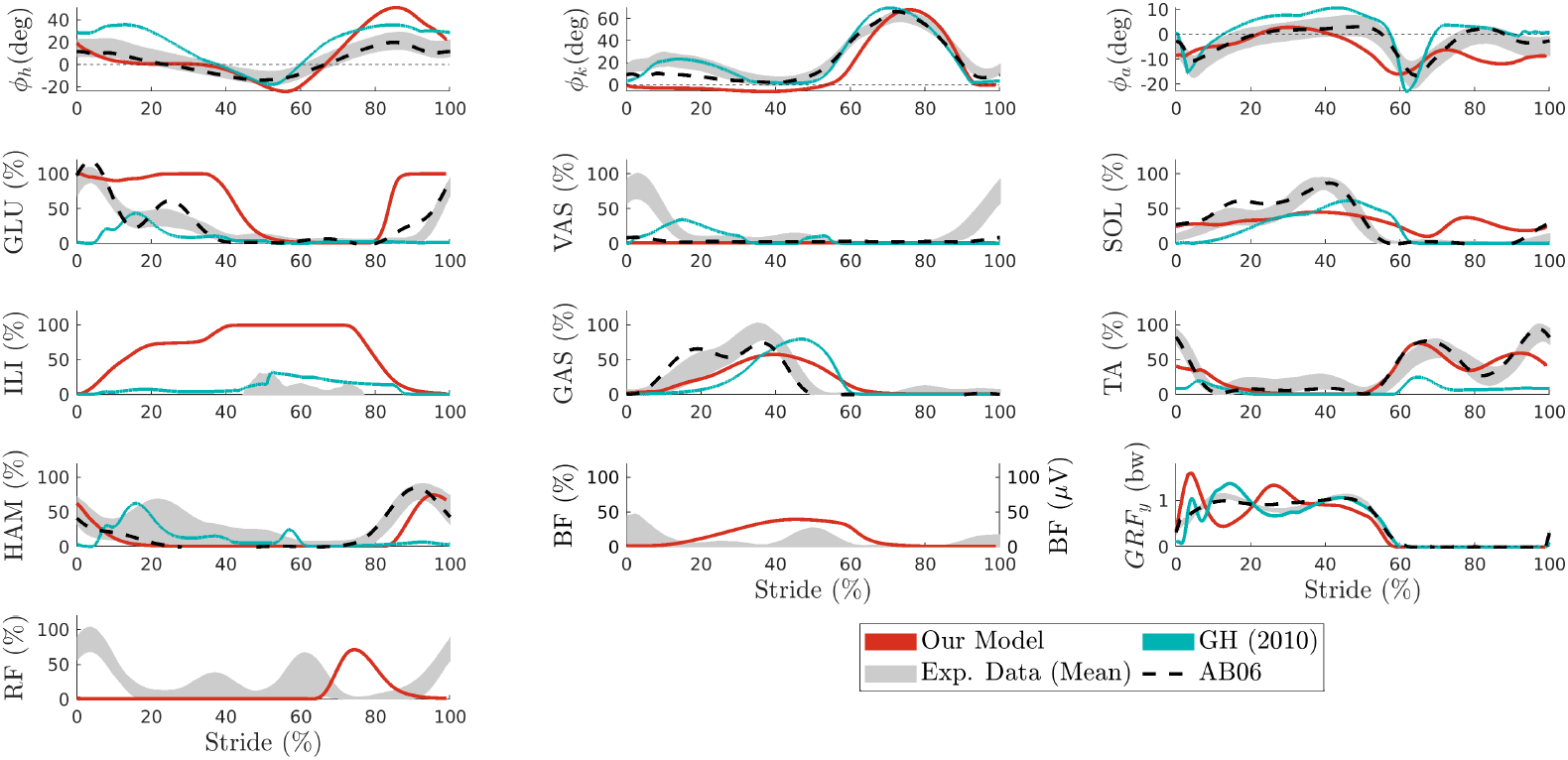
Forwards walking. Joint kinematics (first row) and corresponding muscular activation for hip (left), knee (middle) and ankle (right) joint, as well as ground reaction force (bottom right). Data are shown for the mean of all strides of our forward walking solution (red) compared to the model of [21] (blue), experimental data averaged over all subjects from [51, 53, 54] (grey area, see also Section “Experimental data”) and of one height and weight matched subject AB06 from [51] (dotted black).

Also in terms of muscular activations the controller produces smooth muscular activations which match experimental data reasonably well. Maximum cross-correlation values can be found in Table S3. The straight knee during stance is caused by an absence of VAS activation. Among the variations of forwards walking we found, there is also one gait with a knee flexion during stance and VAS activation as well as less shifted ground reaction forces (see Fig. S1b for kinematics and activations and Table S3 and S4 for maximum cross-correlation results). Therefore, this mismatch is likely a result of either the lack of reflex modulation that makes it difficult to differentiate between stance and swing, or the generic cost function, or a combination of both. Note, however, that the gait is still close to human walking as humans exhibit a wide variety of gait kinematics and muscular activations (e.g. height and weight matched subject AB06 from [51] (see Fig. 4 and Table 3) also shows very little VAS activation and a relatively straight leg during stance, *R*_K,AB06-our_ = 0.95).

For our running targets, we discovered various solutions with different running velocities. Fig. 5 shows the kinematics and muscular activations for running gaits at different velocities ranging from 2.72 m/s to 3.42 m/s. While ankle kinematics are not accurately matched (*phi*_a_: *R* = 0.25, Δ = 10 %), hip and knee angle match experimental data well (*ϕ*_h_: *R* = 0.91, *ϕ*_k_: *R* = 0.93) even though as in Walk Fwd the knee is too straight during stance. Also in Run, ground reaction forces are shifted and initially ground reaction forces are substantially higher in the model than in the experimental data (*R* = 0.78, Δ = 8.5 %). Except for VAS, which is not activated at all and RF which misses activation during stance, muscular activations show a fair agreement with experimental data (0.8 *< R <* 0.91, see Table S5), but some muscles saturate at maximum activation (GLU, SOL, ILI, GAS, TA) probably because only speed was maximized and effort was not taken into account.

**Fig 5.**
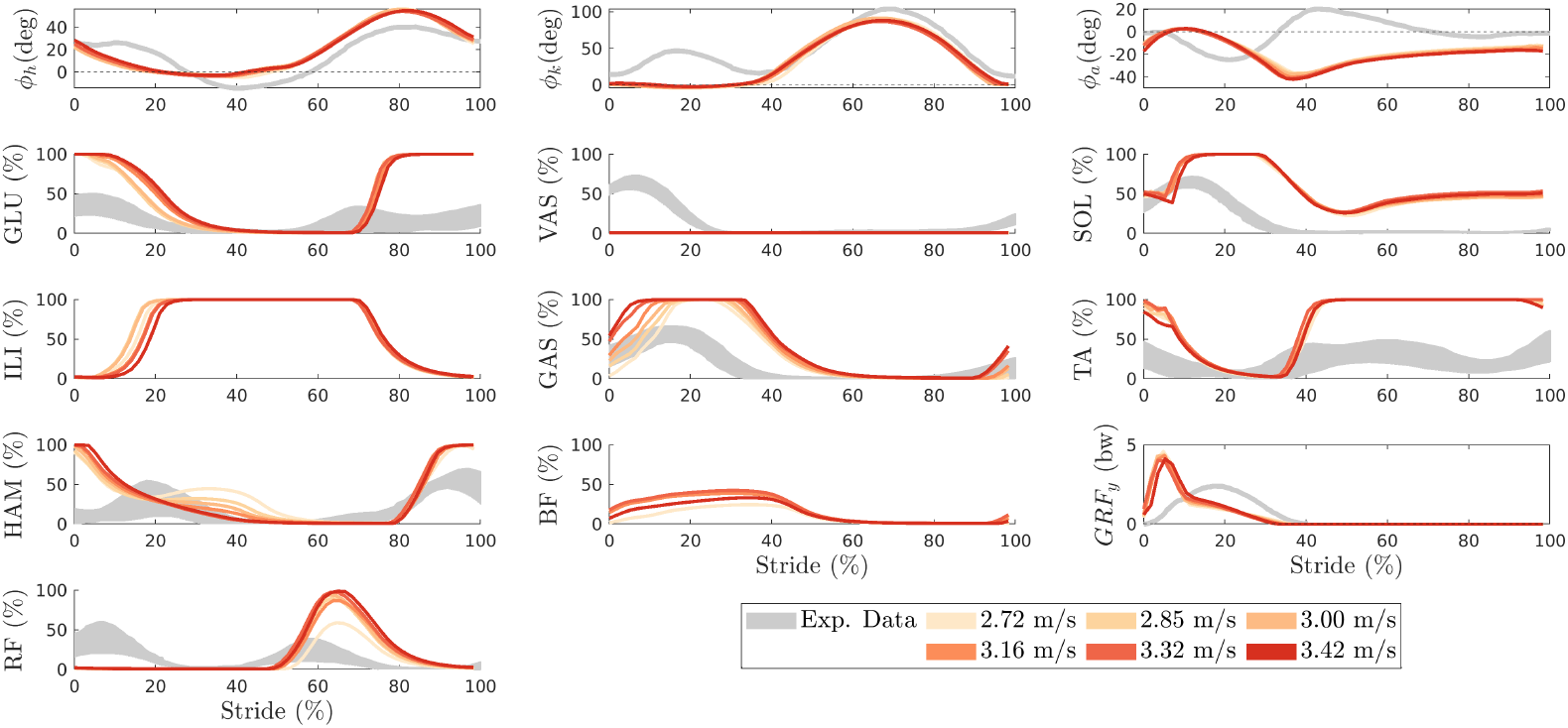
Running. Joint kinematics (first row) and corresponding muscular activations for hip (left), knee (middle) and ankle (right) joint as well as ground reaction force (bottom right) for running at different velocities in comparison to experimental data of running at 3.0 m/s from [55] (grey).

The kinematics and muscle activations for Walk Bkw, Hop Fwd and Hop Bkw are provided in Fig. S4. In an iterative, hand-tuning process we were furthermore able to find a wide spectrum of solutions for each target gait (see Video S2). Also, a variety of other related gaits including skipping or backwards running can be generated. To illustrate the versatility of the reflex controller, we compiled a collection of some other interesting gaits we found (see Video S3). While these gaits are often suboptimal, they still demonstrate a lively appearance that is difficult to quantify.

A comparison of the parameter values of the five target gaits provides some insights into the parameter space of the controller (see Fig. 6). Generally, the solutions are very diverse and differ a lot between the different gaits as well as within gait (see Fig. S2 for parameter values of Walk Fwd 2, a second, manually optimized solution for Walk Fwd). A direct analysis of reflexes is difficult, as the interference between reflexes can cancel out the influence of a reflex such that its value becomes meaningless (e.g. a large negative prestimulation will cause an inactive muscle even if other reflexes would generate an activation of this muscle). However, more generally we find that the offsets *c*_*i*_ are higher for the gaits without effort minimization (hopping and running) than for the walking gaits, where effort minimization was included. Also, homonymous feedback was not only excitatory for the length reflexes, but also for most of the force reflexes, even though this was not a constraint during the optimization. Overall, we find that for all five target gaits, most reflexes are active, with only a few reflex gains set to an absolute value below 0.1 (Walk Fwd: 3, Walk Bkw: 5, Hop Fwd: 1, Hop Bkw: 4, Run: 2). However, as argued before and as the example of VAS during walking and running shows, not all of these reflexes have an influence as they might get canceled out by prestimulations or other reflexes. In fact, the additional walking solution Walk Fwd 2 generates forwards walking with overall smaller absolute reflex gains and 19 reflexes below 0.1(compare also Fig. S2).

**Fig 6.**
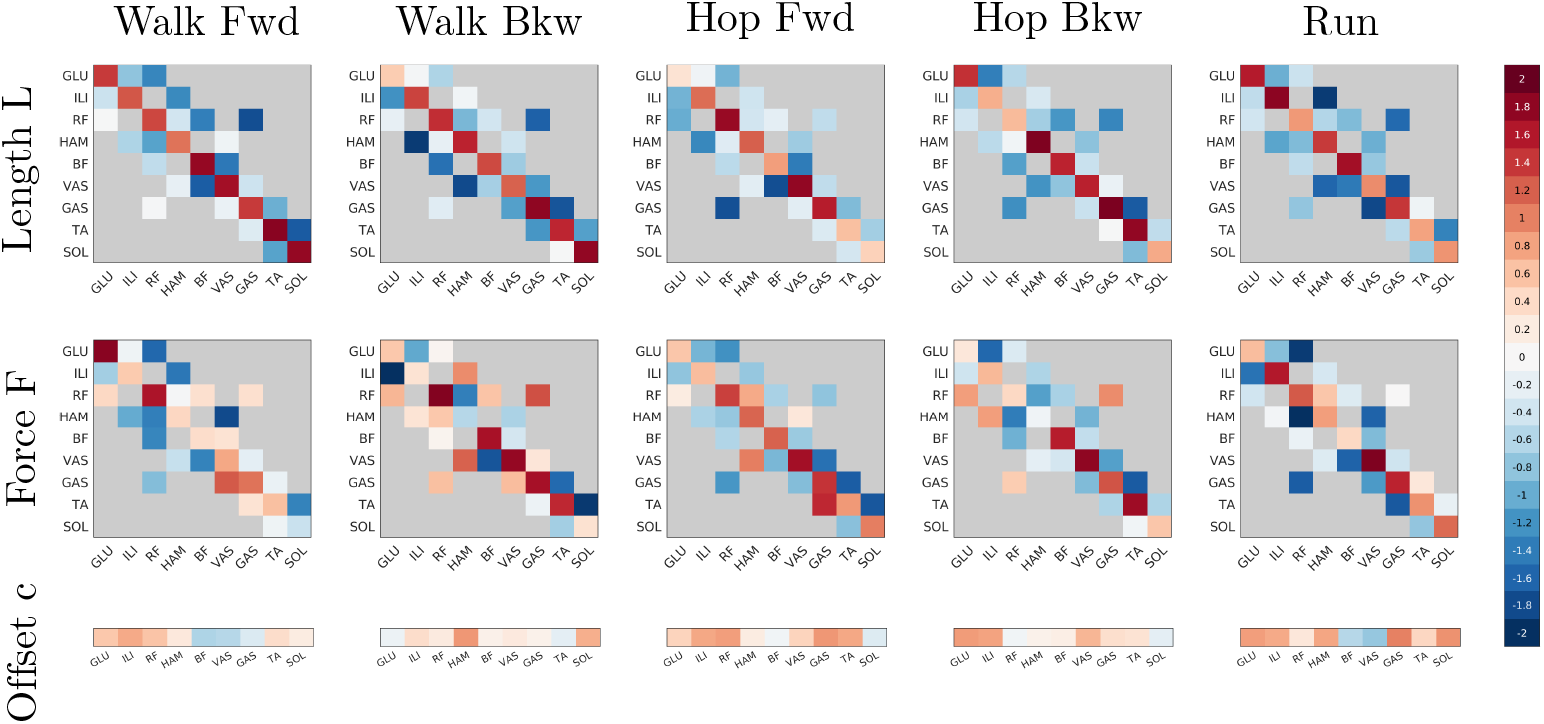
Controller parameters. Parameter values for the five target gaits (left to right) for length (top row) and force (middle row) reflexes as well as the constant prestimulations *c*_*i*_ (bottom row).

## Discussion

Our results demonstrate that, contrary to current beliefs [31], no state-switching mechanism or central pattern generator is required to produce stable bipedal gaits. A simple set of reflexes with constant feedback gains is enough for rhythmic locomotion behavior to emerge.

Since the controller does not depend on hand-picked connections or a finite-state machine specifically tailored to walking, it can furthermore generalize to other natural-looking gaits besides walking, exposing the remarkable potential of reflex-based control. This natural versatility in the absence of any high-level component constitutes one of the main strengths of our controller, which still has to be unveiled to its full extent and is difficult to quantify. Our manually found solutions give a first qualitative impression of the versatility within and between gait, however, future studies should perform a thorough analysis of the parameter space.

The absence of a state switch mechanism and the restriction to homonymous and antagonistic length and force feedback are the main differences of our control architecture compared to the controller of [21]. However, due to its reflexive nature, our controller shares many features with the controller of [21] and subsequent works [22, 26]. First, inter-stride variability is normally relatively low as the periodic movement is based on an interplay of sensor signals. Second, the initial state of the model is important and set close to the target motion as the reflex controller is dependent on the sensory feedback to get into equilibrium. Third, kinematics can not be tracked easily, they can however be used in the cost function to discover reflex parameters that lead to similar kinematics. On the other hand, kinematics emerge naturally from an interplay between body mechanics, muscles and control and generally muscular activations are smooth and natural without abrupt changes.

Our controller has the ability to naturally converge to human-like activations and walking kinematics, without the use of experimental data during the optimization. Kinematically, the main difference to experimental data is the straight knee during stance. The controller proposed by [21] better reproduces knee angle and ground reaction forces (while having less similarity with the ankle angle). However, if we do not compare to the mean kinematics of all subjects but only to one height and weight matched subject, our reflex controller reaches similar kinematic similarity as the controller of [21]. Also, when manually tuning the controller parameters, we can generate forwards walking with a flexed knee during stance and better matching ground reaction forces. Therefore, the mismatch in kinematics is to a large extent caused by the generic cost function and optimization approach. Also, we suspect that reflex modulation might help to properly differentiate between passive VAS during swing and active VAS during stance.

In terms of muscular activation, the results are similar to the ones of the model of [21]: for some muscles our controller better tracks the experimental data, while for others the controller of [21] is more accurate, and both models do show differences to experimental data. Generally, experimental studies show there is a wide range of muscle activity patterns both intra-subject between strides as well as inter-subjects [57, 58]. Similarly, our controller produces different forwards walking solutions with some differences in the kinematics (i.e. knee flexion during stance), but also large differences in muscle activation patterns of several muscles (GLU, VAS, HAM, BF for Walk Fwd compared to Walk Fwd 2). Therefore, it is difficult to make a fair quantitative comparison. The within-gait variability of our reflex controller allows for different patterns, while the whole spectrum that can be covered with the controller still has to be explored.

Purposefully, our controller is as simple as possible, as we tried to include as little reflexes and components as possible to study their potential, before incrementally increasing the complexity of the controller (Occam’s razor). We only use length and force information showing that this is sufficient as sensory input for stable rhythmic gaits. Other sensory information such as muscle velocity, ground contact, leg load, joint angles or vestibular feedback, commonly employed by other works [21, 22] are not required in our controller, keeping the basic control structure simple. While there is no doubt that there are many more sensor signals and components involved in human motor control [5, 7], we follow a bottom-up approach and only include a minimal set of sensor signals. Adding trunk lean or generally vestibular information could be a future addition to offload the stabilization task from the current reflexes and ease stable gait generation. Especially, for a possible extension of the model to 3D the increased difficulty to stabilize the model might require the addition of vestibular feedback like in [21, 22]. Another extension of the controller especially relevant for targeted movements and more complex environments as well as a better kinematic fit of the knee angle could be the addition of reflex gain modulation over the gait cycle [7, 29], which we do not cover in the controller yet.

A further difference to the controller of [21] is the generality of our controller. The controller of [21] has been tailored to level-ground walking and therefore is not so easily extended to other gaits. [22] impressively extended the controller to allow for more complex movements like turning movements and running. However, for this not only reflexes were added but also a supraspinal control layer, while [59] found the trunk lean to be an important variable to modulate running speed. The reflex controller we propose in this work, can generate running (and backwards walking as well as hopping) without any change to the controller architecture, only using length and force reflexes. Furthermore, different running speeds can be obtained by varying the reflex parameters, solely through length and force reflexes. The reached maximum velocity of 3.42 m/s is well below the maximum velocity professional athletes can run at (average velocity for world record for marathon: 5.8 m/s or 5000 m: 6.62 m/s [60]), however it is close to the mean preferred running speed of 3.7 m/s found in experienced runners [61]. Furthermore, an increase of maximal muscle forces (mimicking training) could likely increase the maximum velocity of our model to a certain extent as well and also the simplicity of the model and lack of arms might contribute to the slower speed [62].

All presented results are the starting point for further development and studies to overcome its current limitations. We adopted the current approach to allow studying the broad versatility of the control structure. When focusing on one specific gait the differences to experimental data could possibly be minimized by a more complex cost function including further optimization criteria like joint pain or kinematic tracking. Kinematic tracking could also provide a way to model subject differences and analyze differences in the resulting reflexes. Even though it might seem counter-intuitive in a forward simulation, for reflex controllers this provides a neat way to constrain the kinematics as the reflex controller can not directly follow a trajectory and therefore the kinematics only shape the cost function.

In the same line of studying the broad versatility of the controller we allowed for all homonymous and antagonistic reflexes such that the potential of the controller could be explored and no bias was introduced by selecting reflexes more specifically. This, however, limits the analysis and interpretability of the found solutions, as likely not all reflexes are needed for all gaits and interaction effects between parameters can lead to reflexes without effect, even if the reflex gain is high. In the future these limitations could be overcome by a detailed study of the parameter space including a minimization of active reflexes, an analysis of the connections within as well as between gaits, and an assessment of the initial state dependency. With this, the dependency on the cost function and hyperparameters could be eliminated providing a more continuous view on the parameter space. A minimization of reflexes could help to obtain a sparser controller structure and a more meaningful set of reflexes that can better be compared to neurophysiological data and reduces the number of redundant or unnecessary reflexes. The focus of this work was to showcase the versatility and potential of the control structure, however the additional solution for forwards walking we found by manual tuning and iterative optimization already demonstrates that potentially less reflexes are necessary and might lead to more natural-looking results.

Long-term the controller promises a valuable basis for the study of gait modulation (e.g. speed) and transitions by analyzing the connections between parameter sets. Also, currently, a change of gait means a complete change of parameters which is not very plausible to happen in humans, even though reflexes do change e.g. between postural control and locomotion [63]. However, all these further studies together might shed more light on the common structure allowing for an even more reduced basic control structure that includes the versatility and simplicity shown in this work while allowing a neurological interpretation and gait transitions in a biologically plausible way. Also, the proposed controller provides a valuable basis for extensions to more complex and targeted, voluntary movements as well as different terrains and perturbations demanding a higher-level control structure.

Overall, our proposed methodology is sufficient to demonstrate the power of the spinal reflex control approach without high-level state selection or modulation. To our knowledge, this is the first work that allows to generate different gaits solely based on reflexes without any higher-level control structure, finite-state machine or CPG. The controller cannot yet reproduce human locomotion completely, and aspects such as gait transitions or goal directed movements are not integrated yet. However, it provides a solid starting point to study the relation between different gaits and develop more complex neuromuscular controllers up to the integration of voluntary control [30]. Several works have shown the potential of reflex controllers in the control of exoskeletons, prosthesis and robots [13, 14, 64]. Due to its independence from gait state detection the controller could ease its application in the control of these systems and its versatility could furthermore allow to extend their functionalities.

## Conclusion

Our results demonstrate that proprioceptive reflexes are a remarkably powerful control primitive and that state-switching with a finite state machine or a CPG is not needed to generate rhythmic movement. This suggests that simple proprioceptive reflexes are sufficient to produce the basic rhythmic activity required for human locomotion. Furthermore, our reflex controller allows for a striking versatility of gaits suggesting that spinal reflexes play an even more prominent role than expected. Even though we know that other circuits, such as central pattern generators, play an important role in gait, their importance, relation and dependencies might need to be re-evaluated. Our proposed controller provides a valuable basis to develop more complex neuromuscular controllers and has possible applications in the control of robots, exoskeletons, prostheses or ortheses.

## Supporting information

Upon publication, the controller, model and optimization results will be made available open source.

## Acknowledgments

E.K.B. and S.S. acknowledge the support by the Stuttgart Center for Simulation Science (SimTech). E.K.B. was furthermore supported by the International Max Planck Research School for Intelligent Systems and the German Academic Scholarship Foundation.

## Author contributions

Conceptualization, E.K.B., S.S. and T.G.; data curation, E.K.B. and T.G.; formal analysis, E.K.B. and T.G.; investigation, E.K.B. and T.G.; methodology, E.K.B., D.F.B.H, S.S. and T.G.; project administration, E.K.B., S.S.; software, E.K.B. and T.G.; validation, E.K.B.; visualization, E.K.B.; writing–original draft, E.K.B. and T.G.; writing–review & editing, E.K.B., D.F.B.H, S.S. and T.G.; resources, S.S.; funding acquisition, S.S.; supervision, D.F.B.H, S.S. and T.G.;

