## Supplementary material for "A simple network of proprioceptive reflexes can produce a variety of bipedal gaits"

### Model

**Table S 1. Model parameters.** Optimal fiber length $l_{opt}$, tendon slack length $l_{slack}$, maximum isometric force $F_{max}$, pennation at optimal fiber length $\alpha$ (taken from [1], with exception of hamstring parameters which were taken from [2]), muscle grouping and modelled delays $\Delta t$ (taken from [3], homonymous and antagonistic reflex, for antagonistic reflexes the mean of both involved muscles is used) of the 9 Hill-type muscles per leg. For more details see Section "Model" of main publication.

| | $l_{opt}$(cm) | $l_{slack}$(cm) | $F_{max}$ (N) | $\alpha$ (°) | Muscles | $\Delta t$ (ms) Hom. | Ant. |
|---|---|---|---|---|---|---|---|
| GLU | 15.69 | 11.1 | 1944 | 21.9 | Gluteus max. super. + middle + infer. | 10 | 10 |
| ILI | 10.66 | 15.2 | 2186 | 14.3 | Psoas | 10 | 10 |
| RF | 7.59 | 34.49 | 1169 | 13.9 | Rectus fem. | 20 | 20 |
| HAM | 9.76 | 31.9 | 2594 | 11.6 | Biceps fem. l. head, semimem. semiten. | 15 | 20 |
| BF | 11.03 | 9.5 | 804 | 12.3 | Biceps fem. s. head | 20 | 20 |
| VAS | 9.93 | 12.31 | 4530 | 4.5 | Vastus intermedius, lateralis, medialis | 20 | 20 |
| GAS | 5.1 | 38.4 | 2241 | 9.9 | Gastrocnemius lateral + medial head | 35 | 35 |
| TA | 6.83 | 24.3 | 1579 | 9.6 | Tibialis anterior | 35 | 35 |
| SOL | 4.4 | 24.8 | 3549 | 28.3 | Soleus | 35 | 35 |

**Table S 2. Initial model state.** Initial state values $\phi$ (rad or m) and velocities $\dot{\phi}$ (rad/s or m/s) of the model for all five target gaits, manually tuned and obtained from the gait cycle of initially found solutions for the movement.

| | Walk Fwd | | Walk Bkw | | Run | | Hop Fwd | | Hop Bkw | |
|---|---|---|---|---|---|---|---|---|---|---|
| | $\phi$ | $\dot{\phi}$ | $\phi$ | $\dot{\phi}$ | $\phi$ | $\dot{\phi}$ | $\phi$ | $\dot{\phi}$ | $\phi$ | $\dot{\phi}$ |
| pelvis_tilt | -0.03 | -0.57 | -0.1 | -0.05 | -0.08 | 1.39 | -0.4 | 1.68 | -0.3 | 0 |
| pelvis_tx | 0 | 1 | 0 | -0.8 | 0 | 4.49 | 0 | 1.55 | 0 | -0.5 |
| pelvis_ty | 0.95 | 0.05 | 0.95 | -0.03 | 0.94 | -0.63 | 0.91 | 0.72 | 0.8 | 0 |
| hip_flexion_r | 0.17 | -0.58 | 0.1 | -0.3 | -0.19 | 2.11 | 0.6 | -6.74 | 0.7 | 0 |
| knee_angle_r | -0.1 | -1 | -0.1 | -0.05 | -0.85 | -9.72 | -0.36 | 5.77 | -1 | 0 |
| ankle_angle_r | 0.02 | 0.99 | 0.2 | 0.1 | -0.78 | 0.46 | 0.16 | -1.9 | 0.5 | 0 |
| hip_flexion_l | 0.23 | 4.91 | 0.7 | -3 | 0.24 | -6.87 | 0.6 | -6.74 | 0.7 | 0 |
| knee_angle_l | -1.23 | -8.6 | -1.1 | -0.3 | 0.06 | -0.03 | -0.36 | 5.77 | -1 | 0 |
| ankle_angle_l | -0.17 | 0.63 | -0.2 | -0.1 | -0.11 | 7.4 | 0.16 | -1.9 | 0.5 | 0 |

### Target gaits

#### Walk Fwd

**Table S 3. Maximum cross-correlation $R$ and time shift $\Delta$ (%) for muscular activation** Comparison of mean of experimental data from [4–6] (exp) or mean data of subject AB06 from [4] and simulation data from our model (our) or the model of Geyer & Herr [7] (GH) as well as the second forwards walking solution Walk Fwd 2 (our,2).

| | Glu | | Vas | | Sol | | Ili | | Gas | |
|---|---|---|---|---|---|---|---|---|---|---|
| | $R$ | $\Delta$ | $R$ | $\Delta$ | $R$ | $\Delta$ | $R$ | $\Delta$ | $R$ | $\Delta$ |
| exp-our | 0.81 | 0 | 0.58 | 0 | 0.82 | 1.8 | 0.69 | 3.7 | 0.97 | 7.4 |
| exp-GH | 0.86 | 10.4 | 0.74 | 9.7 | 0.99 | 8.8 | 0.88 | 4.7 | 0.99 | 11.2 |
| exp-our,2 | 0.72 | 20 | 0.55 | 1.2 | 0.85 | 1.1 | 0.64 | 13.5 | 0.73 | 0 |
| AB06-our | 0.83 | 0 | 0.91 | 0 | 0.83 | 0.6 | – | – | 0.97 | 11.5 |
| AB06-GH | 0.83 | 10.8 | 0.65 | 8.8 | 0.98 | 9.7 | – | – | 0.94 | 14.3 |

| | Ta | | Ham | | Bf | | Rf | |
|---|---|---|---|---|---|---|---|---|
| | $R$ | $\Delta$ | $R$ | $\Delta$ | $R$ | $\Delta$ | $R$ | $\Delta$ |
| exp-our | 0.91 | 6.0 | 0.83 | 0 | 0.55 | 9 | 0.37 | 14 |
| exp-GH | 0.82 | 0 | 0.55 | 0 | – | – | – | – |
| exp-our,2 | 0.93 | 0 | 0.94 | 0 | 0.8 | 0 | 0.33 | 8.8 |
| S06-our | 0.94 | 0.3 | 0.86 | 1.2 | – | – | – | – |
| S06-GH | 0.83 | 0 | 0.35 | 6.4 | – | – | – | – |

**Table S 4. Maximum cross-correlation $R$ for joint angles and GRF** Similarity metrics computed between mean of experimental walking data (exp) and simulation data of the second forwards walking solution of our model (our,2) in comparison to the results of our target gait Walk Fwd (our) and the model of [7] (GH) already given in the main results Tablel 3.

| | $\phi_h$ | | $\phi_k$ | | $\phi_a$ | | GRF | |
|---|---|---|---|---|---|---|---|---|
| | $R$ | $\Delta$ | $R$ | $\Delta$ | $R$ | $\Delta$ | $R$ | $\Delta$ |
| exp-our,2 | 0.84 | 1.3 | 0.97 | 0.7 | 0.76 | 3.9 | 0.97 | 2.3 |
| exp-our | 0.90 | 0 | 0.89 | 2.1 | 0.79 | 3.4 | 0.93 | 3.6 |
| exp -GH | 0.93 | 0 | 0.98 | 0.6 | 0.69 | 1 | 0.99 | 0.7 |

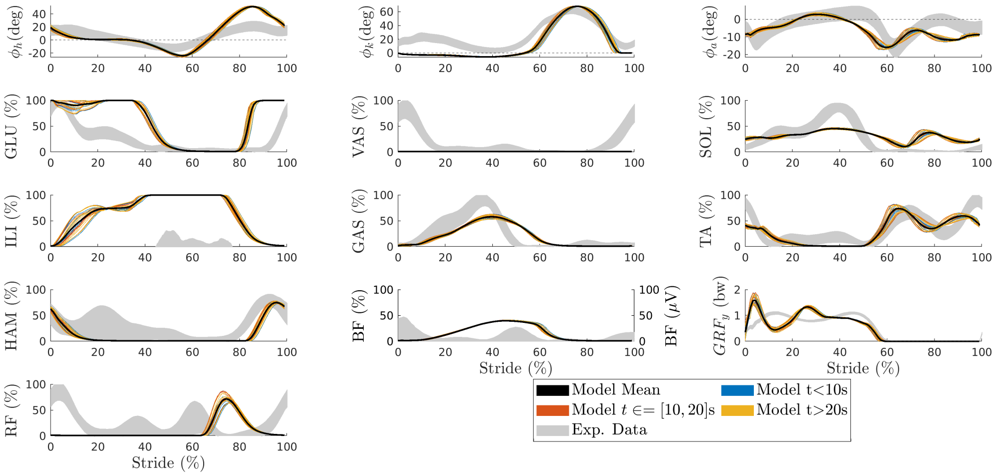


**(a)** Walk Fwd ($1.2\,\mathrm{m\,s^{-1}}$)

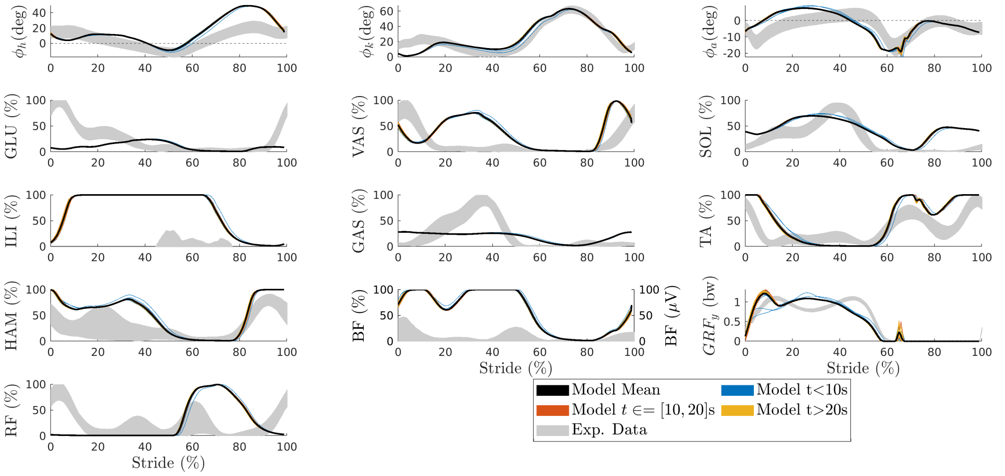


**(b)** Walk Fwd 2 ($1.1\,\mathrm{m\,s^{-1}}$)

**Fig S 1. Kinematics, GRF and muscular activation** Joint angles and muscular activations for hip, knee and ankle joint (left to right) as well as ground reaction force for (a) our target gait Walk Fwd and for (b) a different parameter set Walk Fwd 2 generating forwards walking with knee flexion during stance and VAS activation. Strides are colored based on the time interval they start in ($t < 10\,\mathrm{s}$, $t \in 10\,\mathrm{s}$ to $20\,\mathrm{s}$, $t > 20\,\mathrm{s}$) and maximum simulation time is set to $t_{\max} = 30\,\mathrm{s}$.

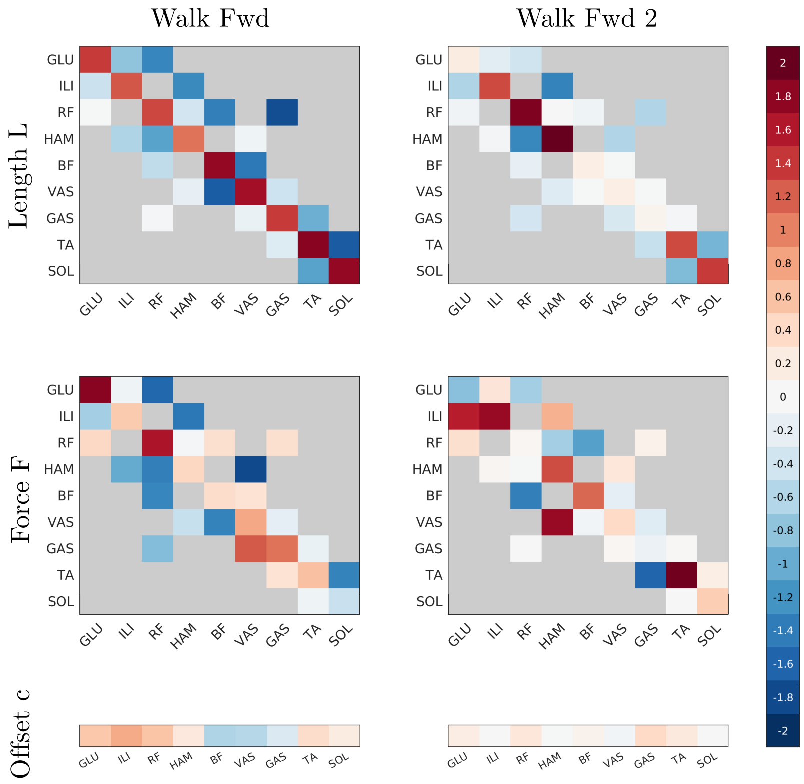


**Fig S 2. Parameter sets Walk Fwd** Comparison of two parameters sets generating forwards walking: Walk Fwd (left) the found target gait and Walk Fwd 2 (right) that was manually found and exhibits more knee flexion during stance.

#### Run

**Table S 5. Maximum cross-correlation $R$ and time shift $\Delta$ (%) for muscular activation during Run** Computed between mean of experimental data (exp) from [8] and simulation data from our model (our).

| | Glu | | Vas | | Sol | | Ili | | Gas | |
|---|---|---|---|---|---|---|---|---|---|---|
| | $R$ | $\Delta$ | $R$ | $\Delta$ | $R$ | $\Delta$ | $R$ | $\Delta$ | $R$ | $\Delta$ |
| exp-our | 0.89 | 0 | 0.56 | 0 | 0.82 | 8.6 | – | – | 0.91 | 2.8 |

| | Ta | | Ham | | Bf | | Rf | |
|---|---|---|---|---|---|---|---|---|
| | $R$ | $\Delta$ | $R$ | $\Delta$ | $R$ | $\Delta$ | $R$ | $\Delta$ |
| exp-our | 0.95 | 0 | 0.85 | 0 | – | – | 0.56 | 7.7 |

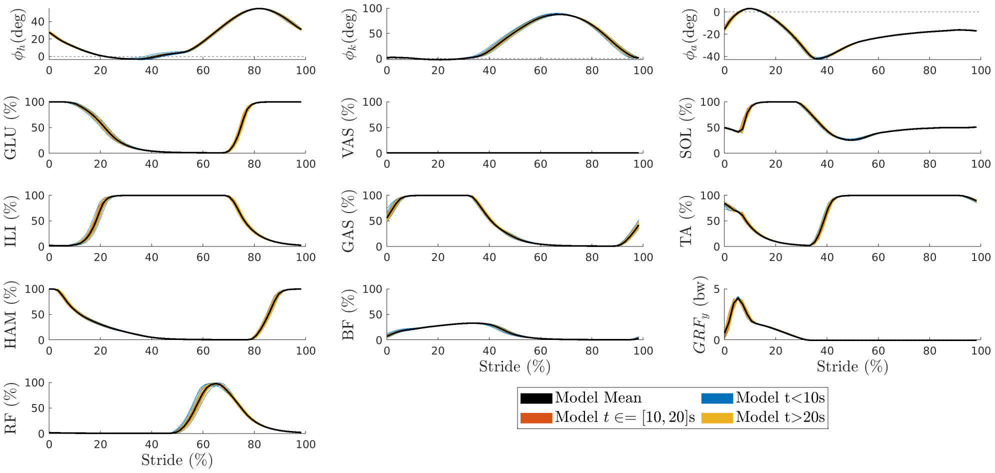


**Fig S 3. Kinematics, GRF and muscular activation** Joint angles and muscular activations for hip, knee and ankle joint (left to right) as well as ground reaction force for Run. Strides are colored based on the time interval they start in ($t < 10$ s, $t \in 10$ s to 20 s, $t > 20$ s) and maximum simulation time is set to $t_{\max} = 30$ s.

#### Other target gaits

A) Walk Bkw at 1.21 m/s

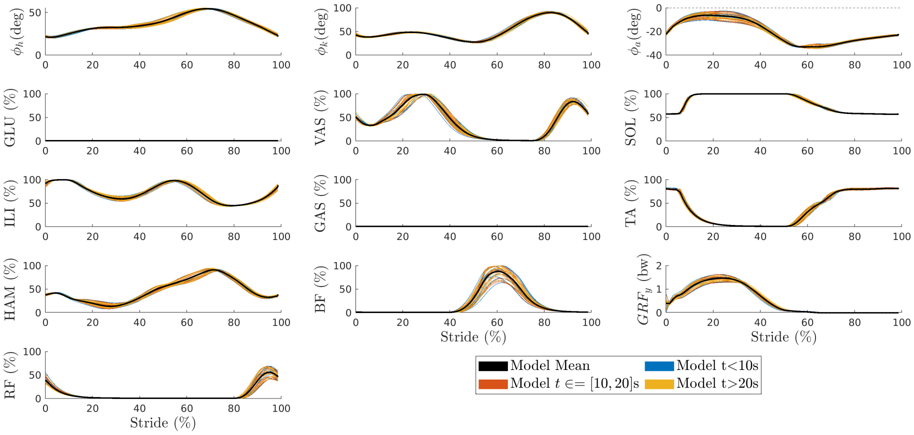


B) Hop Fwd at 1.22 m/s

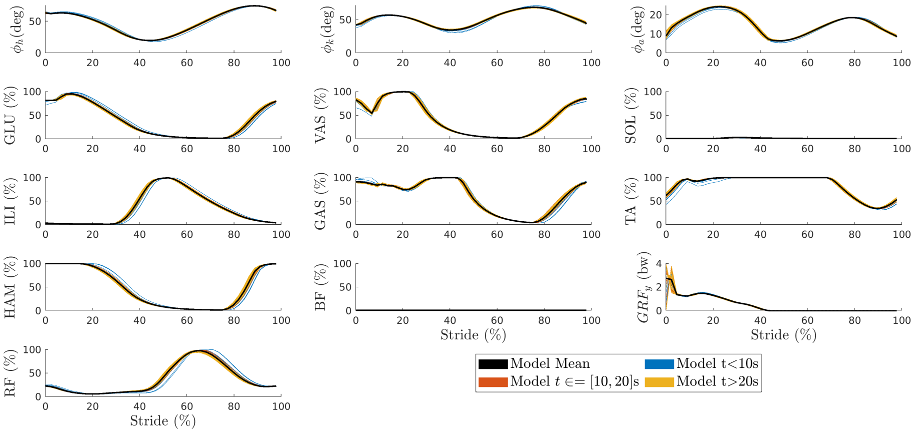


C) Hop Bkw at 1.39 m/s

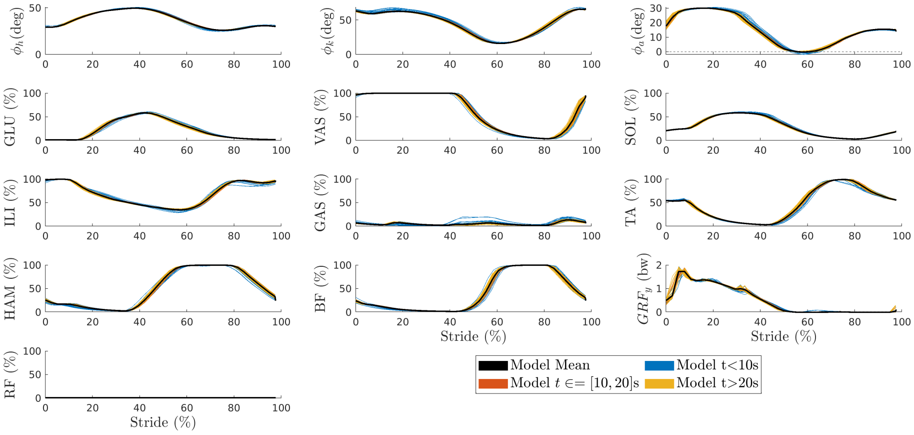


**Fig S 4. Kinematics, GRF and muscular activation** Joint angles and muscular activations for hip, knee and ankle joint (left to right) as well as ground reaction force. Strides are colored based on the time interval they start in ($t < 10\,\mathrm{s}$, $t \in 10\,\mathrm{s}$ to $20\,\mathrm{s}$, $t > 20\,\mathrm{s}$) and maximum simulation time is set to $t_{\max} = 30\,\mathrm{s}$.
